# Speed-dependent modulation of tactile edge orientation discrimination

**DOI:** 10.1101/2025.09.24.678395

**Authors:** Vaishnavi Sukumar, J. Andrew Pruszynski

**Author notes:** **Corresponding Author** J. Andrew Pruszynski, Robarts Research Institute, Room 1232H, Western University, London, Ontario, Canada, N6A 5B7, E. **Author Contributions** Vaishnavi Sukumar : Conceptualization, Data acquisition, Formal analysis, Investigation, Methodology, Visualization, Writing - original draft and editing; J. Andrew Pruszynski : Conceptualization, Supervision, Funding acquisition, Investigation, Visualization, Methodology, and Writing - review and editing. **Data and materials availability** All raw data generated in this study can be made available upon request. **Competing Interest** The authors declare no competing interests.

## Abstract

Previous studies investigating edge-orientation discrimination capacity either stimulated an immobilized finger at limited speeds, or did not manipulate movement speed during active exploration. Here we tested the effect of movement speed on edge-orientation discrimination, including very slow and fast speeds. Participants were instructed to move their finger across two pairs of edges while matching their speed to a visual cue. One edge pair was parallel and the other non parallel to varying degrees. Participants were asked to identify the non parallel pair of edges. We report three main findings. First, consistent with previous reports, when they were free to choose their movement speed, participants moved at an average speed of ∼29 mm/s (range: 15-52 mm/s). Second, there was no correlation between a participant’s natural speed and their edge orientation discrimination capacity. Third, participants got better at edge orientation discrimination at slower than average speeds (5mm/s), and worse at higher than average speeds (90 and 180 mm/s). This change in performance was correlated with their relative change in movement speed.

**New and Noteworthy:** Here we demonstrate speed-dependent variation in edge-orientation discrimination during active tactile exploration, with improved performance at very slow speeds. Changes in discrimination capacity as a function of speed are correlated with deviations from a participant’s natural speed.

## Introduction

One important aspect of object manipulation is the ability to decipher shape and form using tactile cues, like the orientation of an object relative to the fingertips. For instance, to successfully button a shirt, one needs to be able to grasp the button, orient the button’s edge to align it with the buttonhole and then place the button through the hole while keeping it aligned. Since the button is generally occluded by the fingers, this behavior must arise because of the information signaled by first-order tactile neurons, likely those innervating low threshold mechanoreceptors in the glabrous skin of the fingertips (Johansson & Flanagan, 2009). Indeed, digital anesthesia applied locally to the fingertips impairs performance in such behaviors that require precision object control (Johansson et al., 1992; Monzée et al., 2003).

We previously showed that single first-order tactile neurons signal edge orientation differences (Pruszynski & Johansson, 2014; Sukumar et al., 2022) that are at least as small as 5° apart for a range of stimulation speeds (15-180mm/s). The peripheral neural signaling of edge orientation differences is better at stimulation speeds slower (15-20 mm/s) than the average behavioral speed of ∼30 mm/s and falls off sharply at higher speeds (Sukumar et al., 2022). Although previous studies have shown that people use a range of speeds during tactile behavior in general (Callier et al., 2015; Cole & Abbs, 1988; Johansson & Westling, 1984; Olczak et al., 2018; Smith, Chapman, et al., 2002; Smith, Gosselin, et al., 2002), Vega-Bermudez and colleagues were the first to show that faster stimulus presentation speed leads to a decrease in tactile acuity (Vega-Bermudez et al., 1991). However, the effect of slower speeds on tactile perception of fine details like edge orientation differences is unknown.

In this study, we explored the effects of slow (5 mm/s), average (30 mm/s) and fast (90 mm/s and 180 mm/s) movement speeds on tactile acuity. Our goal was to establish the relationship between movement speed and edge orientation discrimination capacity. We used a wide range of edge orientation differences (4-20°), centered on the average edge discrimination acuity previously reported using the same apparatus (∼12°; Olczak et al., 2018). We report three main findings. First, consistent with previous reports, our participants moved at an average speed of ∼29 mm/s (range: 15-52 mm/s) (Olczak et al., 2018; Vega-Bermudez et al., 1991). Second, also consistent with our previous work, there was no correlation between a participant’s natural speed and their edge orientation discrimination capacity (Olczak et al., 2018). Third, similar to the signaling capacity of first-order tactile neurons (Sukumar et al., 2022), participants got better at edge orientation discrimination at slower than average speeds (5mm/s), and worse at higher than average speeds (90 and 180 mm/s). This change in performance was correlated with the change in movement speed relative to an individual’s natural speed.

## Methods

### Participants

A total of 28 young adults (age range : 20 - 32 years, average age : 25) volunteered for the study. Participants reported no known conditions that affected their tactile acuity or hand control. All participants provided written, informed consent to participate in the study, and were compensated for their participation. The study was approved by the Western University Human Research Ethics Board, and was performed in accordance with the Declaration of Helsinki.

### Apparatus

Participants sat in front of a screen that prevented them from seeing the edge stimuli (Figure 1A). Participants extended their index finger, and a reflective marker (Figure 1B) was taped on the nail of the index finger without occluding the finger pad. Behind the screen were two magnetic plate holders that held two sets of plates. We used plates similar to the ones previously described (Olczak et al., 2018). Briefly, the plates were 6 cm wide and were covered by an acrylic slot that limited the region of the plate that could be touched (white area on the plate in Figure 1B) and kept the contact surface constant. Each plate was covered in a nylon material (Toyobo EF 70 GB), which contained pairs of raised edges. Each edge was 5 mm high, 8 mm wide at its base and 5 mm wide at its peak. The first edge in every plate was always perpendicular to the direction of motion. The second edge was either parallel to the first edge (parallel, 0° edge orientation difference), or angled away from the direction of motion to varying degrees (non-parallel, 4°, 8°, 12°, 16° or 20° edge orientation differences). For both parallel and non parallel plates, the distance between the edges at their midpoints was always 15 mm.

**Figure 1.**
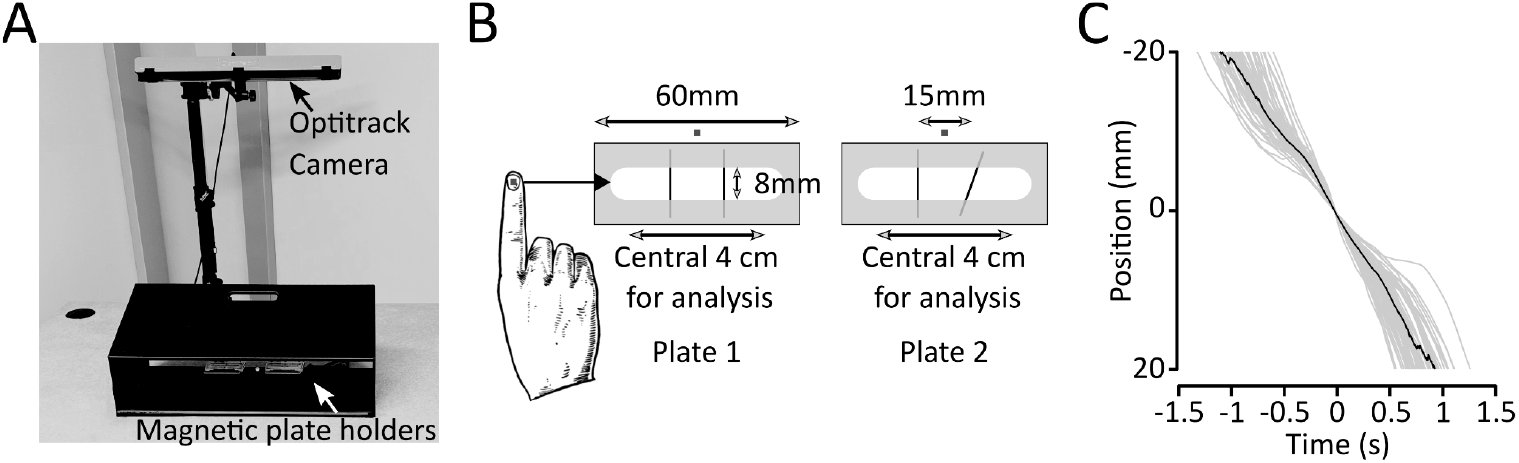
Experimental setup. A) Edge plates were set in magnetic plate holders that were placed inside a box that occluded participants’ vision. B) A pair of representative plates, one with parallel and the other with non parallel edges. Grey squares on the fingernail and in front of the plates represent reflective markers for the OptiTrack cameras. C) Sample kinematic traces from an exemplary participants, across 60 trials (gray) and their median (black).

### Data Acquisition

A motion capture system (V120:Trio, OptiTrack, NaturalPoint, Inc.) tracked a reflective marker placed on the participant’s fingernail. Permanently fixed markers were also used to mark the center of the plates and the apparatus, which was kept constant between sessions. A pre-calibrated tool was used to set the origin of the three dimensional space being recorded. The output from the system provided the three-dimensional coordinates of all markers used.

### Experimental Design

The experiment was based on a two-alternative forced choice protocol. In each trial, the participant placed the fingertip of their dominant hand in the acrylic slot (white area on the plate in Figure 1B) and moved it over the edges in each plate in succession. The plates were switched after every trial. Participants were asked to verbally report which plate had the non-parallel edge, and were provided with verbal feedback about whether they were correct. The location of the parallel and non-parallel plates were fully randomized. The edge orientation difference for the non-parallel plate was also fully randomized. We also randomly used 5 different parallel plates so that participants could not base their decision on some idiosyncratic imperfection in the parallel plate.

Our experiment included six blocks, with each block having sixty trials. In block 1, participants passed their index finger over the edges and reported non-parallel edges, similar to our previous work (Olczak et al., 2018). Each edge orientation difference (4°, 8°, 12°, 16° or 20°) was presented twelve times, for a total of sixty trials. Participants were not instructed about how quickly to move their finger relative to the stimulus. In the remaining five blocks, participants were asked to follow a visual cursor and match its speed while exploring the edges. The speed of this visual cursor varied - 5mm/s, 30mm/s, 90mm/s, 180mm/s, or the participant’s natural speed calculated from block 1. The slowest speed, 5mm/s, was picked based on a pilot study where participants were asked to move as slow as they could while moving their fingers over the plates with the raised edges. Before each block, participants were allowed to practice with the apparatus, and speed stimuli for the later blocks, so as to build familiarity with the setup. We used two plates where edges were parallel (0° edge orientation difference). These two plates were not reused during the experiment outside of the practice trials. The practice sessions, lasting approximately five minutes each time, were not analysed.

### Inclusion Criteria

All volunteers were able to reliably discriminate our largest edge difference (20°). In order to study the effect of the speed, participants had to be able to reliably discriminate at least 16° edge orientation differences at their natural speed, so that a decrease in performance with changing speed could be measured. Reliable discrimination was defined as accurately identifying the edge orientation difference at least 75% of the time. That is, of the 12 trials presented for the 16° edge orientation differences, participants had to correctly report the non-parallel plates in at least 9 trials. Six participants were not able to meet this criteria. One participant was unable to understand the instructions and could not complete the task. One participant was unable to follow the speed cues and could not complete the task. The remaining participants (N = 20) are used in all the analyses.

### Data Analysis

Data was analyzed using custom analysis scripts (MATLAB, 2023a, MathWorks, Natick, MA). Statistical analyses were performed in GraphPad (GraphPad Prism version 10.0.0 for Windows, GraphPad Software, Boston, MA).

## Results

### Natural speed does not predict discrimination capacity

Twenty participants completed six blocks, performing 360 trials each (5 edge differences x 12 trials per edge difference = 60 trials per block, six blocks). In the first block, participants performed the task at a self-chosen (i.e. natural) speed. Figure 1B shows the plates mounted on magnetic holders. The central 4 cm of the plates, surrounding the embossed edge, was used to calculate the average finger speed. Figure 1C illustrates the finger movement of an exemplar participant in the first block, where they were free to choose their movement speed, i.e., their natural speed. This participant moved consistently across the plate with an average natural speed of ∼20mm/s. Natural speed ranged from 14-58 mm/s across individuals, with a population average speed of ∼29 mm/s (Figure 2A). Speed was not significantly different between the plates (paired t-test, t(19) = 1.93, p = 0.07). Participants maintained relatively constant speed throughout the first block (Figure 2C). Although we observed a trend suggesting participants adapted their speed early in each block, this was not statistically significant (paired t-test between average velocity in the first and last five trials : t(19) = 1.43, p = 0.17).

**Figure 2.**
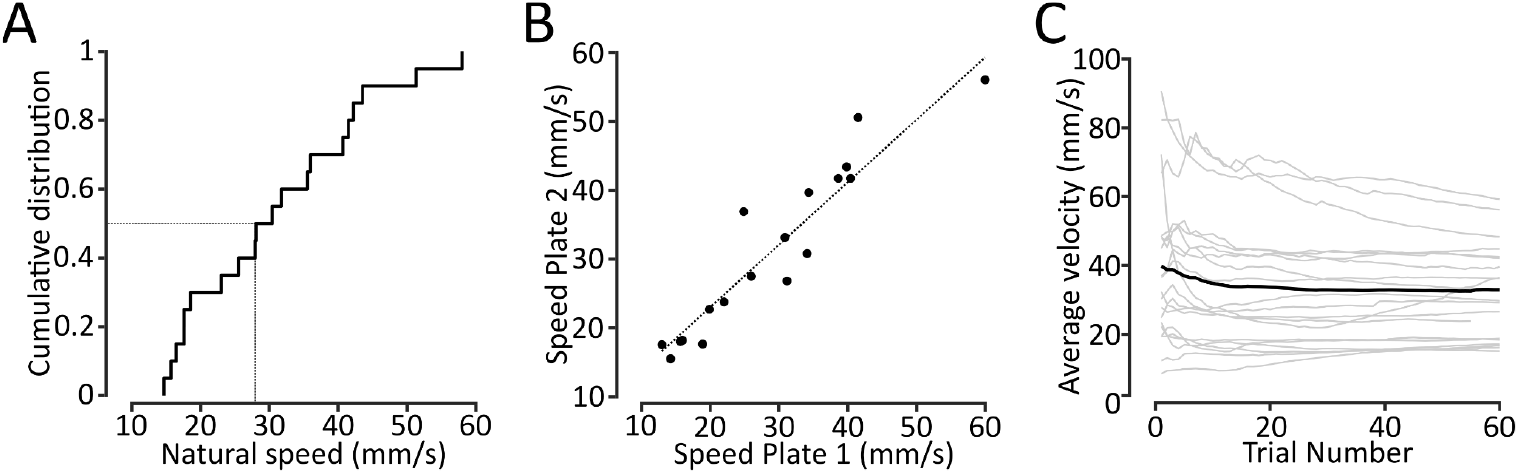
Natural speeds. A) Average movement speeds for each participant in the first block, where participants moved at their natural speed. B) Average natural speed for each participant across both the plates. C) Average natural speed for each trial in the first block, for each participant (grey lines). Black line shows the population average.

Participants moved at similar speeds across both plates, regardless of whether the edges were parallel or non parallel, and irrespective of trial accuracy (Figure 3A). A two-way repeated measures ANOVA applied to the median participant speed across correct and incorrect trials over parallel and non parallel plates showed no effect of plate condition (F_1,19_ = 2.56, p = 0.126) or accuracy (F_1,19_ = 0.86, p = 0.365). Interestingly, despite the broad range of natural speeds (14 - 58 mm/s, Figure 2A), we found no reliable relationship between a participant’s natural speed and their discrimination capacity (Figure 3B, r^2^ = 0.0001 for overall performance, r^2^ < 0.11 for each edge difference). That is, a participant’s natural speed was not predictive of their edge orientation discrimination capacity.

**Figure 3.**
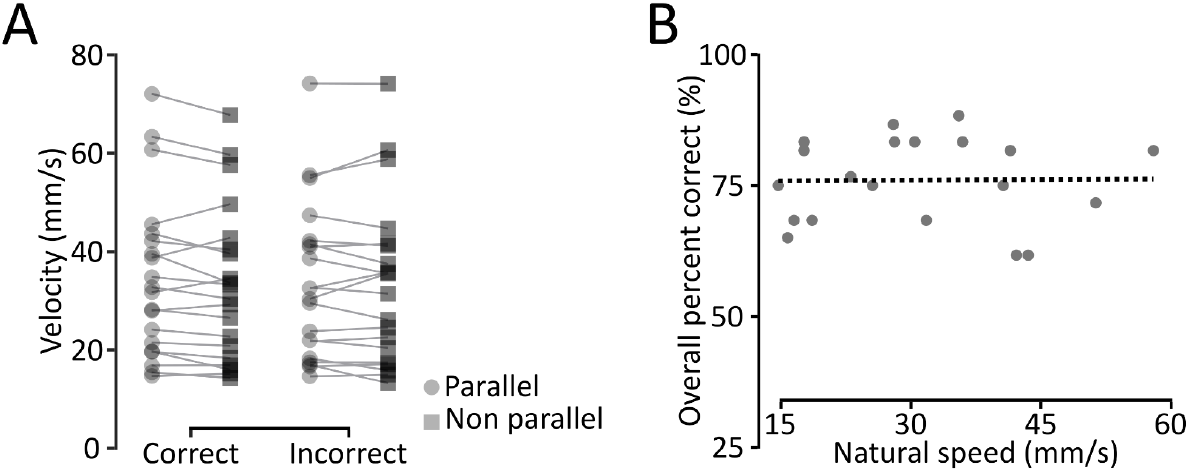
Natural speed does not reflect discrimination capacity. A) Average natural speed across parallel and non parallel plates for each participant in the first block, for both correct and incorrect trials. B) Participant performance as a function of their natural speed. The dashed line shows the linear regression fit.

### Manipulating speed affects discrimination capacity

In blocks 2-6, participants were asked to match their movement speed to visual cues moving at 5, 30, 90 or 180 mm/s in four of the blocks. One of the blocks featured a visual cue that moved at the participant’s natural speed (as measured in the first block). All five cued speed blocks were randomized across participants. Participants exhibited higher variance in speed matching at the faster speeds (90 and 180 mm/s; Figure 4A; r^2^ = 0.69), and were on average slower than the cue at these faster speeds. In contrast, participants matched slower speeds more accurately. Despite the dual nature of the visually cued blocks, we found no change in edge orientation discrimination capacity between the first block (where participants moved without any speed constraints at their natural speed) and in the subsequent block where they were cued at their natural speed (paired t-test between average velocities in each block across participants, t(19) = 1.421, p = 0.172; Figure 4B). That is, the visual cue we used to manipulate movement speed did not appear to affect discrimination capacity, at least at the natural speed.

**Figure 4.**
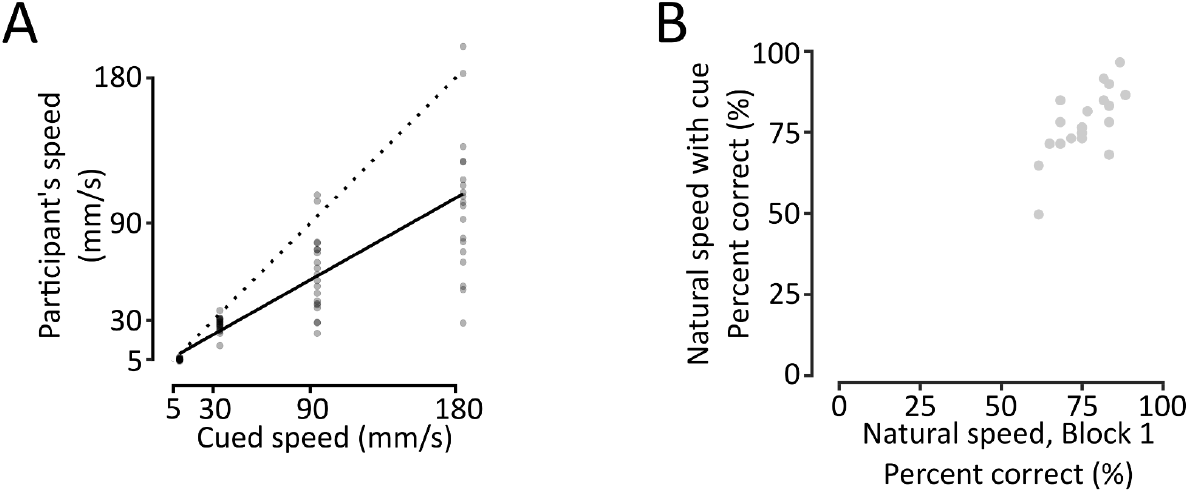
Participants’ response to the cued speeds. A) Relationship between participants’ actual speed in response to the visually cued speed. The dotted line shows the line of unity. The solid black line shows the regression line (r^2^ = 0.69). Most participants were able to match the cued speeds well at slower speeds but not at the faster cued speeds. B) Relationship between the participant’s natural speed (without cues) and their response with visual cues representing their own natural speed.

We next asked if edge orientation discrimination capacity was modulated by the movement speed across the cued blocks including slow (5 mm/s), average (30mm/s) and very fast (90 and 180 mm/s) speed conditions. Figure 5 shows the participants’ overall discrimination capacity across these blocks. Consistent with previous reports (Vega-Bermudez et al., 1991), faster movement speeds led to a decrease in edge orientation discrimination capacity (one-way repeated measures ANOVA, F_3,57_ = 7.723, p < 0.001). We next examined if the effect of speed differed for the smallest and largest edge differences (4° vs. 20°). As expected, our analysis revealed that tactile discrimination capacity decreased with increasing speed (effect of speed: two-way repeated measures ANOVA, F_3,57_ = 10.16, p < 0.0001) for both edge differences (edge effect: two-way repeated measures ANOVA, F_1,19_ = 248.9, p < 0.001). Moreover, the change in discrimination capacity was modulated by the edge difference (interaction effect: F_3,57_ = 2.99, p < 0.05). Decomposing the interaction revealed that the improvement in performance seen between the slowest speed (5 mm/s) and the average speed (30 mm/s), was driven by the finest edge orientation difference (4°). In contrast, for the 20° edge difference, discrimination accuracy remained unchanged between the 5 and 30 mm/s conditions, likely due to a ceiling effect, as performance was already at its maximum for both speeds.

**Figure 5.**
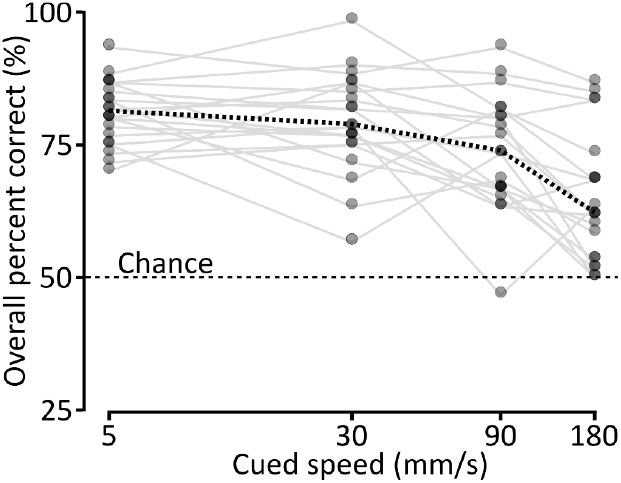
Tactile discrimination as a function of cued speed. The overall percentage of correct responses for each cued speed block. Thin gray lines show individual participants. The dotted black line shows the population median.

To better understand this edge-dependent effect for the slow speed, we compared participants’ discrimination capacity at their natural speeds with their discrimination capacity at the slowest cued speed (5mm/s) for each edge difference (4°, 8°, 12°, 16° and 20°). We found that participants performed better at the slow speed, and this was mainly driven by their improved performance for the smallest (4°) edge difference (Figure 6). A two-way repeated measures ANOVA across the five edge-orientation differences and two speed conditions showed an effect of edge-orientation difference (F_4,76_ = 83.04, p < 0.0001) as well as an effect of speed (F_1,19_ = 5.437, p = 0.031), but no interaction. Post-hoc paired t-tests revealed a significant difference in performance only for the 4° edge condition (t(19) = 2.897, p = 0.009; Figure 6). That is, participants performed better when they moved slower than their natural speeds, and this improvement reflected their performance on the finest edge differences.

**Figure 6.**
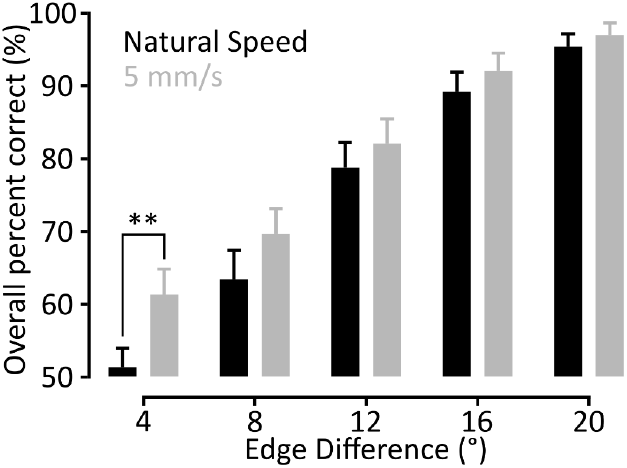
Comparing the performance at slowest speed condition with the performance at the natural speed condition from block 1. Comparisons are shown for each edge pair. Moving slower increased the number of correct responses significantly, especially for the finest edge pair (4°).

### Discrimination capacity scales with relative speed

As shown earlier, despite a broad range of natural movement speeds among participants (14 - 58 mm/s, Figure 2A), natural movement speed was not predictive of edge orientation discrimination capacity (Figure 3B). Specifically, although moving slower (5 mm/s) improved edge orientation discrimination, participants who naturally moved slowly (∼15 mm/s) did not outperform those who naturally moved quickly (∼60 mm/s). Therefore, we next asked if the effect of speed on a participant’s discrimination capacity depends on how much the cued speeds deviate from their natural speeds. That is, whether participants’ discrimination capacity is influenced by relative movement speed.

We compared the relative speed of each participant in the four cued blocks (5mm/s, 30mm/s, 90mm/s and 180mm/s) to their relative discrimination capacity. Relative speed was defined as the difference between their speed in the cued block and their natural speed in the first block. Similarly, relative discrimination accuracy was defined as the difference between the percentage of correct responses in the cued block and the first block. Figure 7 illustrates a clear relationship between relative speed and relative discrimination capacity for these four blocks. Indeed, linear regression analysis showed that relative discrimination capacity decreases with increase in relative speed (r^2^ = 0.39, F_(1,78)_ = 49.68, p < 0.0001). Thus, participants’ performance improved systematically as they moved slower relative to their own natural speed.

**Figure 7.**
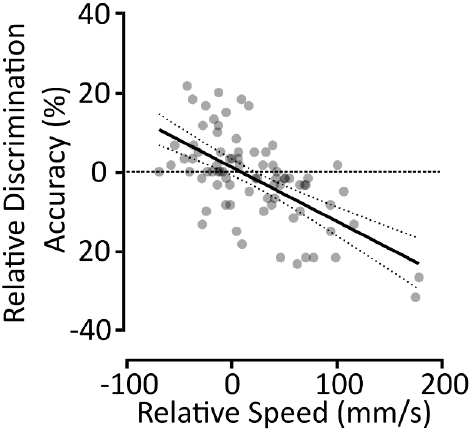
Tactile discrimination as a function of relative speed. Relative discrimination capacity compared as a function of relative speed. Relative speed and discrimination capacity were calculated as the difference between the measures in the cued block and the first block where the participants used their natural speeds in the absence of cues. The solid line shows the regression line, and the dotted lines show the 95% confidence interval.

## Discussion

We investigated the effect of movement speed on edge discrimination capacity. We found that people performed worse for fast speeds compared to slow speeds. This improvement at slow speeds was driven by better performance for finer edge orientation differences (4°). Interestingly, participants had a wide range of natural movement speeds but this did not relate to differences in their edge orientation discrimination capacity (Figure 3B). At the same time, edge orientation discrimination capacity was tightly linked to their own natural speed. That is, people performed better or worse when moving slower or faster compared to their natural speed.

### Edge Orientation Discrimination: Neural Mechanisms and the Action - Perception Gap

Edge orientation discrimination relies on inputs from multiple classes of first-order tactile neurons innervating low threshold mechanoreceptors in the glabrous skin (Johansson & Flanagan, 2009). Fast-Adapting Type-1 (FA-1) neurons, which are sensitive to dynamic skin deformation, respond primarily at the onset and offset of contact with an object. In contrast, Slow-Adapting Type-1 (SA-1) neurons fire during sustained indentation and are active throughout the contact with the object. Previous work has shown that both FA-1 and SA-1 neurons signal large edge orientation differences with comparable fidelity (≥22.5°, Pruszynski & Johansson, 2014), but for finer discriminations, FA-1 neurons outperform SA-1 neurons especially at slow speeds (5°, Sukumar et al., 2022).

Although first-order tactile neurons can signal differences in edge orientation that are as fine as 5°, perceptual discrimination thresholds reported in previous studies are considerably higher. Specifically, when edge orientation stimuli are passively applied to the distal finger pads, participants typically require orientation differences of 15-30° to make reliable discriminations (Bensmaia et al., 2008; Lechelt, 1992; Peters et al., 2015). Unlike texture perception, where active exploration is known to improve tactile acuity (Hollins et al., 2002; Hollins & Risner, 2000), we previously demonstrated that edge orientation discrimination did not improve substantially with active exploration (threshold ∼12°, Olczak et al., 2018). However, when individuals use tactile information to guide object manipulation, edge orientation acuity can be remarkably fine, on the order of ∼3° (Pruszynski et al., 2018).

One proposed explanation for this discrepancy in discrimination acuity between object manipulation and perceptual reporting is that central processing of sensory information is task and context-dependent (Crapse & Sommer, 2008; Engel et al., 2001; Gazzaley & Nobre, 2012; Manita et al., 2015; Schroeder et al., 2010; Zagha et al., 2013). Indeed, this idea is paralleled in the visual system, where sensory input can be uniquely processed for motor and perceptual tasks (Bridgeman, 2000; Kravitz et al., 2011; Milner & Goodale, 2006; Weiskrantz, 1996). Moreover, in touch, motor-related signals can interact with the tactile signaling pathway (Adams et al., 2013; Canedo, 1997; Fanselow & Nicolelis, 1999; Lee et al., 2008; Manita et al., 2015; Seki & Fetz, 2012; Zagha et al., 2013), and thereby influence context-dependent processing of tactile signals such that the same edge orientation information is processed differently for the perceptual and motor tasks.

Our study suggests an additional explanation for this action-perception gap in edge orientation discrimination capacity. Our results suggest that moving slower than one’s natural speed can improve discrimination accuracy. This raises the possibility that previous perceptual studies may have underestimated tactile acuity as they did not explore the relationship between movement speed and tactile acuity at the slower speeds. That is, a thorough estimate of edge orientation discrimination thresholds at slow speeds (e.g., 5 mm/s) may narrow the discrepancy between perceptual thresholds (∼12°, Olczak et al., 2018) and that observed during object manipulation (∼3°, Pruszynski et al., 2018).

### Tactile speed

Previous studies reported that people moved at ∼20 mm/s speed when discriminating fine details like letters and edge orientations for perceptual tasks (Olczak et al., 2018; Vega-Bermudez et al., 1991). In our previous work, similar to our current work, we also found that people moved at a range of speeds, and this natural variation across people had no clear relationship to their tactile acuity (Olczak et al., 2018). Vega-Bermudez and colleagues (1991) were among the first to confirm that increasing the speed decreased tactile acuity for fine form, in their case a letter recognition task. That is, people’s variation in natural speeds has no impact on their tactile acuity, but moving very fast detrimentally affects their tactile discrimination capacity. In our current study, we add to this in two ways. First, we show that moving very slowly, at ∼5 mm/s, significantly improves tactile discrimination, especially for the finest edges. Despite the added benefit, people likely don’t use the slowest speeds naturally, perhaps in a bid for time efficiency. Second, we show that the effect of speed on tactile discrimination capacity is relative to each individual’s natural speed. That is, participants performed better than their natural speeds when moving slower than their natural speeds, and vice versa (Figure 7).

Interestingly, although we see an improvement in tactile acuity with slow speeds (5mm/s), previous studies have featured edge stimuli that were passively indented onto the finger, i.e. at zero speed, with no improvement in tactile acuity. In one study, Bensmaia et al (2008) compared tactile orientation discrimination between scanned and indented bars. They showed that both conditions had similar results - that is, they did not demonstrate a benefit of the zero speed condition. This suggests that although moving slower could confer some advantages, indentation (i.e. zero speed) would lose this advantage. This absence of improvement seen in zero speed could be attributed to the fact that static stimuli cross fewer receptive fields, and therefore recruit fewer neurons at the population level. At the single neuron level, a static stimulus would fail to elicit the spatiotemporal patterning seen by the sequential stimulation of subfields within a complex receptive field (Pruszynski & Johansson, 2014, Hay & Pruszynski, 2020; Zhao et al., 2018). On the other hand, a moving edge stimulus would not only sequentially stimulate the highly sensitive subfields in a receptive field, it would also stimulate more neurons at the population level. Something similar is observed when very small edges (2 mm long) that span a small portion of the finger pad are used. Orientation acuity becomes worse, going from ∼15° to 90° during perceptual tasks (Peters et al., 2015), and from ∼3° to 11° during object manipulation (Pruszynski et al., 2018). That is, shorter edges that stimulate fewer neurons across the finger pad are harder to perceive. By extension, very slowly moving edges that approach zero speed likely recruit fewer neurons at the population level, and therefore are harder to perceive.

Another important factor to consider is the role of skin mechanics and finger geometry in shaping tactile discrimination. Reduced conformance of the skin to a stimulus because of increased skin stiffness and reduced softness - due to aging, dehydration and even different finger sizes - has been linked to decreased tactile acuity (Deflorio et al., 2023; Li & Gerling, 2023; Peters et al., 2009; Vega-Bermudez & Johnson, 2004). Indeed, softening the skin with hyaluronic acid improves tactile acuity (Li & Gerling, 2023). In the indentation (zero-speed) condition, limited skin deformation and reduced contact area may result in poor conformance to the edge stimuli, thereby limiting discrimination capacity. This is further illustrated in Bensmaia et al (2008), where the authors used a stimulus that consisted of densely packed individual pins that were sequentially activated to mimic a scanning motion with raised edges (Killebrew et al., 2006). Although the pins stimulated the entirety of the distal fingerpad in a sequential pattern, they did not induce the same skin deformations that occur with a true scanning motion as seen in active exploration. The sequentially indented pin arrays yielded poorer orientation thresholds (∼20°) compared to active scanning (∼15°; Olczak et al., 2018). This suggests that the dynamic mechanical interaction between the stimulus and the skin during movement may be essential for enhancing tactile acuity. Taken together, these findings indicate that the absence of meaningful skin deformation at near-zero speeds likely limits both the recruitment of tactile afferents and the formation of spatiotemporal patterns necessary for precise discrimination. This aligns with our current results, which show improved acuity at slow but nonzero speeds, and supports the idea that both neural and mechanical factors contribute to tactile acuity.

## Acknowledgements

This work was supported by a CIHR Project Grant to J.A.P (PJT-186177) and the Canada First Research Excellence Fund (via BrainsCAN). J.A.P. received a salary award from the Canada Research Chairs program.

## References

1. Adams, R. A., Shipp, S., & Friston, K. J. (2013). Predictions not commands: Active inference in the motor system. Brain Structure and Function, 218(3), 611–643. 10.1007/s00429-012-0475-5

2. Bensmaia, S. J., Hsiao, S. S., Denchev, P. V., Killebrew, J. H., & Craig, J. C. (2008). The tactile perception of stimulus orientation. Somatosensory & Motor Research, 25(1), 49–59. 10.1080/08990220701830662

3. Bridgeman, B. (2000). Interactions between vision for perception and vision for behavior. In Beyond Dissociation: Interaction Between Dissociated Implicit and Explicit Processing (Vol. 22, pp. 17–40). Amsterdam ; Philadelphia, PA : J. Benjamins.

4. Callier, T., Saal, H. P., Davis-Berg, E. C., & Bensmaia, S. J. (2015). Kinematics of unconstrained tactile texture exploration. Journal of Neurophysiology, 113(7), 3013–3020. 10.1152/jn.00703.2014

5. Canedo, A. (1997). Primary Motor Cortex Influences On The Descending And Ascending Systems. Progress in Neurobiology, 51(3), 287–335. 10.1016/S0301-0082(96)00058-5

6. Cole, K. J., & Abbs, J. H. (1988). Grip force adjustments evoked by load force perturbations of a grasped object. Journal of Neurophysiology, 60(4), 1513–1522. 10.1152/jn.1988.60.4.1513

7. Crapse, T. B., & Sommer, M. A. (2008). Corollary discharge across the animal kingdom. Nature Reviews Neuroscience, 9(8), 587–600. 10.1038/nrn2457

8. Deflorio, D., Di Luca, M., & Wing, A. M. (2023). Skin properties and afferent density in the deterioration of tactile spatial acuity with age. The Journal of Physiology, 601(3), 517–533. 10.1113/JP283174

9. Engel, A. K., Fries, P., & Singer, W. (2001). Dynamic predictions: Oscillations and synchrony in top–down processing. Nature Reviews Neuroscience, 2(10), 704–716. 10.1038/35094565

10. Fanselow, E. E., & Nicolelis, M. A. L. (1999). Behavioral Modulation of Tactile Responses in the Rat Somatosensory System. The Journal of Neuroscience, 19(17), 7603–7616. 10.1523/JNEUROSCI.19-17-07603.1999

11. Gallace, A., & Spence, C. (2009). The cognitive and neural correlates of tactile memory. Psychological Bulletin, 135(3), 380–406. 10.1037/a0015325

12. Gazzaley, A., & Nobre, A. C. (2012). Top-down modulation: Bridging selective attention and working memory. Trends in Cognitive Sciences, 16(2), 129–135. 10.1016/j.tics.2011.11.014

13. Hay, E., & Pruszynski, J. A. (2020). Orientation processing by synaptic integration across first-order tactile neurons. PLoS Computational Biology, 16(12), e1008303. 10.1371/journal.pcbi.1008303

14. Hollins, M., Bensmaiä, S. J., & Roy, E. A. (2002). Vibrotaction and texture perception. Behavioural Brain Research, 135(1), 51–56. 10.1016/S0166-4328(02)00154-7

15. Hollins, M., & Risner, S. R. (2000). Evidence for the duplex theory of tactile texture perception. Perception & Psychophysics, 62(4), 695–705. 10.3758/BF03206916

16. Johansson, R. S., & Flanagan, J. R. (2009). Coding and use of tactile signals from the fingertips in object manipulation tasks. Nature Reviews Neuroscience, 10(5), 345–359. 10.1038/nrn2621

17. Johansson, R. S., Hger, C., & Bäckström, L. (1992). Somatosensory control of precision grip during unpredictable pulling loads. III. Impairments during digital anesthesia. Experimental Brain Research, 89(1), 204–213. 10.1007/BF00229017

18. Johansson, R. S., & Westling, G. (1984). Roles of glabrous skin receptors and sensorimotor memory in automatic control of precision grip when lifting rougher or more slippery objects. Experimental Brain Research, 56(3), 550–564. 10.1007/BF00237997

19. Kravitz, D. J., Saleem, K. S., Baker, C. I., & Mishkin, M. (2011). A new neural framework for visuospatial processing. Nature Reviews Neuroscience, 12(4), 217–230. 10.1038/nrn3008

20. Lechelt, E. C. (1992). Tactile spatial anisotropy with static stimulation. Bulletin of the Psychonomic Society, 30(2), 140–142. 10.3758/BF03330421

21. Lee, S., Carvell, G. E., & Simons, D. J. (2008). Motor modulation of afferent somatosensory circuits. Nature Neuroscience, 11(12), 1430–1438. 10.1038/nn.2227

22. Li, B., & Gerling, G. J. (2023). An individual’s skin stiffness predicts their tactile discrimination of compliance. The Journal of Physiology, 601(24), 5777–5794. 10.1113/JP285271

23. Manita, S., Suzuki, T., Homma, C., Matsumoto, T., Odagawa, M., Yamada, K., Ota, K., Matsubara, C., Inutsuka, A., Sato, M., Ohkura, M., Yamanaka, A., Yanagawa, Y., Nakai, J., Hayashi, Y., Larkum, M. E., & Murayama, M. (2015). A Top-Down Cortical Circuit for Accurate Sensory Perception. Neuron, 86(5), 1304–1316. 10.1016/j.neuron.2015.05.006

24. Milner, & Goodale. (2006). The Visual Brain in Action (2nd ed.). Oxford ; New York : Oxford University Press.

25. Monzée, J., Lamarre, Y., & Smith, A. M. (2003). The effects of digital anesthesia on force control using a precision grip. Journal of Neurophysiology, 89(2), 672–683. 10.1152/jn.00434.2001

26. Olczak, D., Sukumar, V., & Pruszynski, J. A. (2018). Edge orientation perception during active touch. Journal of Neurophysiology, 120(5), 2423–2429. 10.1152/jn.00280.2018

27. Peters, R. M., Hackeman, E., & Goldreich, D. (2009). Diminutive Digits Discern Delicate Details: Fingertip Size and the Sex Difference in Tactile Spatial Acuity. The Journal of Neuroscience, 29(50), 15756. 10.1523/JNEUROSCI.3684-09.2009

28. Peters, R. M., Staibano, P., & Goldreich, D. (2015). Tactile orientation perception: An ideal observer analysis of human psychophysical performance in relation to macaque area 3b receptive fields. Journal of Neurophysiology, 114(6), 3076–3096. 10.1152/jn.00631.2015

29. Pruszynski, J. A., Flanagan, J. R., & Johansson, R. S. (2018). Fast and accurate edge orientation processing during object manipulation. eLife, 7, e31200. 10.7554/eLife.31200

30. Pruszynski, J. A., & Johansson, R. S. (2014). Edge-orientation processing in first-order tactile neurons. Nature Neuroscience, 17(10), 1404–1409. 10.1038/nn.3804

31. Schroeder, C. E., Wilson, D. A., Radman, T., Scharfman, H., & Lakatos, P. (2010). Dynamics of Active Sensing and perceptual selection. Current Opinion in Neurobiology, 20(2), 172–176. 10.1016/j.conb.2010.02.010

32. Seki, K., & Fetz, E. E. (2012). Gating of Sensory Input at Spinal and Cortical Levels during Preparation and Execution of Voluntary Movement. Journal of Neuroscience, 32(3), 890–902. 10.1523/JNEUROSCI.4958-11.2012

33. Sinclair, R. J., & Burton, H. (1996). Discrimination of vibrotactile frequencies in a delayed pair comparison task. Perception & Psychophysics, 58(5), 680–692. 10.3758/BF03213100

34. Smith, A. M., Chapman, C. E., Deslandes, M., Langlais, J.-S., & Thibodeau, M.-P. (2002). Role of friction and tangential force variation in the subjective scaling of tactile roughness. Experimental Brain Research, 144(2), 211–223. 10.1007/s00221-002-1015-y

35. Smith, A. M., Gosselin, G., & Houde, B. (2002). Deployment of fingertip forces in tactile exploration. Experimental Brain Research, 147(2), 209–218. 10.1007/s00221-002-1240-4

36. Sukumar, V., Johansson, R. S., & Pruszynski, J. A. (2022). Precise and stable edge orientation signaling by human first-order tactile neurons. eLife, 11, e81476. 10.7554/eLife.81476

37. Vega-Bermudez, F., Johnson, K. O., & Hsiao, S. S. (1991). Human tactile pattern recognition: Active versus passive touch, velocity effects, and patterns of confusion. Journal of Neurophysiology, 65(3), 531–546. 10.1152/jn.1991.65.3.531

38. Vega-Bermudez, F., & Johnson, K. O. (2004). Fingertip skin conformance accounts, in part, for differences in tactile spatial acuity in young subjects, but not for the decline in spatial acuity with aging. Perception & Psychophysics, 66(1), 60–67. 10.3758/BF03194861

39. Weiskrantz, L. (1996). Blindsight revisited. Current Opinion in Neurobiology, 6(2), 215–220. 10.1016/S0959-4388(96)80075-4

40. Zagha, E., Casale, A. E., Sachdev, R. N. S., McGinley, M. J., & McCormick, D. A. (2013). Motor Cortex Feedback Influences Sensory Processing by Modulating Network State. Neuron, 79(3), 567–578. 10.1016/j.neuron.2013.06.008

41. Zhao, C. W., Daley, M. J., & Pruszynski, J. A. (2018). Neural network models of the tactile system develop first-order units with spatially complex receptive fields. PLOS ONE, 13(6), e0199196. 10.1371/journal.pone.0199196

